# Crisscross multilayering of cell sheets

**DOI:** 10.1101/2021.06.22.449403

**Authors:** Trinish Sarkar, Victor Yashunsky, Louis Brézin, Carles Blanch Mercader, Thibault Aryaksama, Mathilde Lacroix, Thomas Risler, Jean-François Joanny, Pascal Silberzan

## Abstract

Simple hydrostatic skeletons such as the Hydra’s consist of two stacked layers of cells perpendicularly oriented. Although this crisscross architecture can be recapitulated in vitro, little is known on the formation of such multilayers starting from a monolayer. In the present article, we show that bilayering of myoblasts results from the organization and activity of the cells originally in the monolayer which can be described as a contractile active nematic. As expected, most of the +1/2 topological defects that are associated with this nematic order self-propel. However, a subpopulation of these defects remains immobile. Perpendicular bilayering occurs exclusively at these motionless defects. Indeed, cells located at the head of these defects converge toward the (immobile) core and accumulate there until they start migrating on top of the tail of the first layer while the tail cells migrate in the opposite direction under the head cells. Since the cells keep their initial orientations, the two stacked layers end up perpendicularly oriented. This concerted process leading to a bilayer is dependent on the apical secretion of Extra Cellular Matrix (ECM) by the cells. Indeed, we evidence the presence of ECM between the cell layers and at the apical surface of the topmost layer. ECM molecules are oriented in the direction of the cells that produce them, which may guide the migration of the subsequent cell layers on their apical side.

**Significance Statement:** Hydrostatic skeletons such as that of the Hydra consist of two stacked layers of cells perpendicularly oriented whose coordinated contraction allows for complex movements. Such crisscross organization is also observed with myoblasts in culture. Confluent monolayers organize in well-aligned domains between which topological defects position themselves. Although these singularities are generally self-propelled, a fraction of them remains motionless. Perpendicular bilayering occurs exclusively at these particular pinned defects. Cells first accumulate at the head of the defects until they split in two perpendicular layers migrating in an antiparallel way on top of each other. Such a concerted process is highly dependent on the precise organization of the cell-secreted Extra Cellular Matrix (ECM).

## Introduction

In contrast with epithelia that can be cultured in vitro as bidimensional monolayers for long times, most other cell types grown on planar substrates develop in the third dimension after confluence. The property to remain a monolayer can be related to the concept of contact inhibition of proliferation by which an increase of cell density results in an increase of the division time (1–5), or (by generalization) to other regulation mechanisms such as the extrusion and apoptosis of individual cells from the monolayer above a certain cell density (5–7).

Non-epithelial cell types that don’t retain a bidimensional cell sheet organization after confluence develop multicellular 3D structures. They can take the form of aggregates on top of the initial monolayer (for instance neural progenitor cells (NPCs) in vitro (8)), or adopt a stratified structure in which the cells differentiate as the multilayered structure develops (for instance skin epidermis or thymus (9)). Of note, transformed epithelial cells that express an oncogene also develop 3D structures after confluence, often coupled with the down-regulation of cell-cell adhesions in the context of the Epithelial-to-Mesenchymal Transition (EMT) (1, 2, 10–13). Because of the physiological importance of this 2D to 3D transition, strategies to include these 3D aspects have been developed when engineering functional tissues in vitro. These efforts have led to the development of Multilayered Cell Cultures and organs or tumors “on chips” (14–16).

Surprisingly, and despite its importance both in vivo and in vitro in the above-mentioned applications, the mechanism by which mammalian cell monolayers develop into a multilayered structure remains elusive. For instance, one can ask whether this transition is initiated by isolated cells in the monolayer that would then somehow extrude from it and proliferate in such way as to develop a second layer progressively covering the surface of the first one, or if it is a concerted behavior, in which the initial monolayer “splits” into two stacked layers that would then expand by migration and / or proliferation while, possibly, differentiate to organize in a mature stack. Another salient related question relates to the locus of multilayering: does it occur at a random location in the monolayer? Is it related to a particular cell or group of cells slightly different from the others? Or to the local organization of the cells in this initial monolayer?

It is worth noting here that the fate of a cell that has been extruded from a monolayer is both cell type-dependent and context-dependent. For instance, MDCK epithelial cells tend to enter apoptosis and die after having been extruded from a monolayer, generally on the apical side (6, 7), but they remain alive and incorporate into tri-dimensional rims when they are located next to a physical boundary (7). In another example of the impact of physical constraints, elongated cells that orient along a common direction extrude preferentially at singularities in the supracellular organization of the cells (i.e. at topological defects) (8, 17).

Furthermore, it has been shown that the multilayer organization of spindle-shape fibroblast-like cells from chick embryos depends on the tissue type the cells originate from. Indeed, the angle between successive layers varies from uncorrelated (cornea cells), to 0° (parallel alignment, skin cells) or 90° (perpendicular or “crisscross” alignment, heart cells) (18, 19). Of notice, when contracted/relaxed independently, stacked perpendicular muscle layers allow complex functions and exploration of the third dimension as exemplified by hydrostatic skeletons that actuate the hydra (20, 21) or the intestines (22), as well as by muscular hydrostats such as animal tongues (23). As a matter of fact, such principles are used in biomimetic designs of “soft robots” (23).

In the present article, we wish to understand the physical and biological processes responsible for the evolution from a cell monolayer to a crisscross bi- and eventually multi-layer.

The recent theories of out-of-equilibrium active matter (24, 25) provide a very general framework to model biological cells and tissues, allowing to understand some aspects of the mechanical behavior of cell monolayers (26–30). In particular, populations of elongated cells have been successfully described as active nematic phases in which cells have no positional order but share a common orientation (30). These continuum theories have for instance provided a framework to understand inplane spontaneous cell flows in monolayers of myoblasts or retinal cells (31), the critical role of the topological defects associated with such symmetries (32), as well as the emergence of turbulence at low Reynolds number in monolayers of bronchial epithelial cells (28).

In addition to physical effects, it has been shown that the multilayering process is directly related to the production of extracellular matrix (ECM) proteins such as collagen, by the cells themselves (33–35). Therefore, it is expected that physical and biochemical contributions concur to the development of a crisscross bilayer.

More specifically, we study here how spindle-shaped C2C12 murine myoblasts transition from a bidimensional active nematic phase into crisscross multilayers at long times. After confluence, most of the classical +1/2 defects that are characteristic of a contractile active nematic system are mobile. Over time, they pairwise annihilate with −1/2 defects. However, a fraction of these defects remains immobile. We find that bilayers form at these immobile +1/2 defects. The peculiar asymmetric velocity field around these particular +1/2 defects results in an accumulation of cells at the head of the defect. The defect then evolves into a bilayer configuration in which the cells originally at the head of the defect migrate collectively *on top* of the initial monolayer while the cells originally pertaining to the tail migrate *under* the head cells in a ECM-dependent process. Since they keep their initial orientations, the cells of the two stacked layers eventually arrange in a crisscross structure. We show that the onset of this multilayering process is largely controlled by the physics of this contractile active nematic system while the antiparallel movements and the perpendicular orientations of the two stacked layers rely also on the presence and structure of ECM proteins secreted by the cells.

## Results

As muscle cells organize frequently in perpendicular stacked layers in vivo (22), we chose to study the bilayering of C2C12 myoblasts. Sparse C2C12 cells plated on Fibronectin-coated glass proliferated, reached confluence (defining a time t_1L_) and then multilayered, reaching thicknesses up to 30 μm, 5 days after confluence (Supplementary Figure 1). As hypothesized, these cells form striking crisscross bilayers (Figure 1a,b). Being interested in the process by which a cell monolayer develops into two stacked perpendicular cell layers, we monitored the behavior of the cells after they have reached confluence and until they form a crisscross bilayer. We call the time of initiation of the bilayering t_2L_. In the present work, we dissect the formation of a crisscross bilayer in relation with both the local organization of the cells and the production of ECM proteins by the cells themselves.

**Figure 1:**
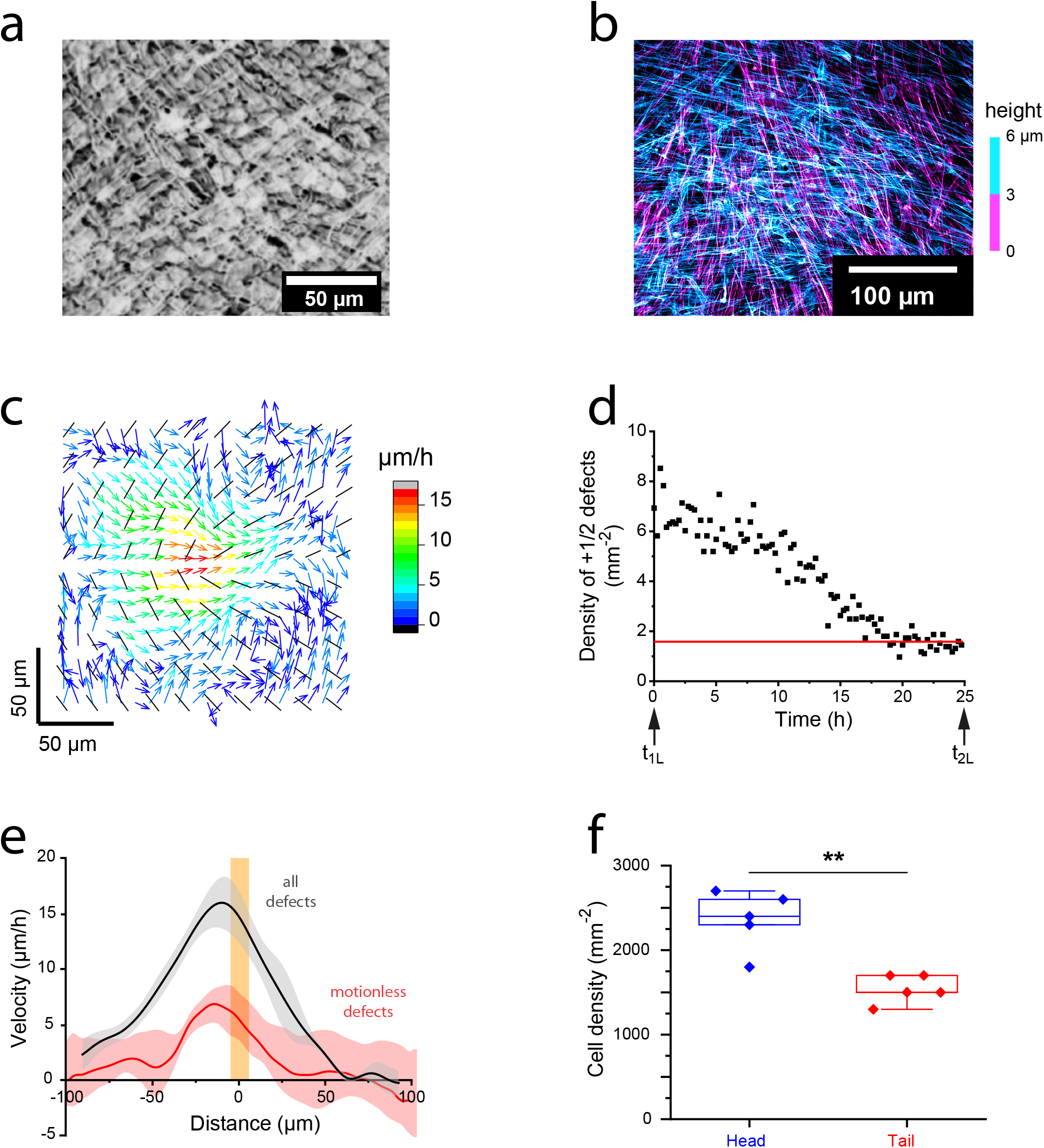
a/ C2C12 bilayer (fixed cells, phase contrast) showing the perpendicular orientation of the two stacked layers. b/ Confocal image of the same showing the two stacked layers at orthogonal orientations (Fixed cells, actin labeling. The color codes for the height.) c/ Before bilayering, the average orientation field (black lines) and the velocity field (colored arrows) measured at a +1/2 defect are consistent with a contractile active nematic description (average over 5398 observations from 350 independent defects). d/ The density of +1/2 defects decreases with time but plateaus at typically 2 defects/mm^2^ several hours before the onset of bilayering (representative experiment). e/ Average velocity profiles along the axis of the +1/2 defects. Black line: all defects (N=350), red line: immobile defects (N=7). The orange bar is the position of the core. Note the zero velocity in the tail of the immobile defects while the velocity in the head remains finite. Colored areas are the SDs. f/ As cells of the head of the immobile defects flow toward their (pinned) core, the cell density increases in the head (measurements within 1h before bilayering).

In the following, we call “layer 1” the first cell layer directly in contact with the coverslip; subsequent layers are called “layer 2”, …, “layer n”.

Note that C2C12 cells are undifferentiated myoblasts that would eventually differentiate into myotubes if cultured in the right conditions of serum and growth factors (36). However, in the present experiments, we use cells that have experienced enough passages after confluence (up to 40) to be selected against differentiability (37, 38). This way, we can study the multilayering phenomenon with cells whose phenotype remains unchanged for the duration of the experiments.

### 1- C2C12 cells organize in an active nematic monolayer

After reaching confluence, C2C12 cells self-organized in a bidimensional active nematic phase in which cells orient along a common direction (30), as previously reported (8, 26, 31, 32). This phase is characterized by typical 2D nematic topological defects (−1/2 and +1/2) that apposition themselves between domains of uniform orientation (Supplementary Figure 2). Such defects can be readily identified in phase-contrast images or fluorescence images of the actin cytoskeleton, and the contractile or extensile nature of the system can be determined from the analysis of the cell flow around +1/2 defects (30). We first analyzed these flows in the 10 hours following confluence. Taking advantage of the characteristic comet-like shape of the +1/2 topological defects, we superimposed a large number (5398 observations from 350 different defects) of them to access their average orientation field and the associated velocity field (39) (Figure 1c). In the present situation, two counterrotating vortices developed at +1/2 defects. Their signs were characteristic of a contractile nematic system (28, 40). Most of the +1/2 defects were therefore motile, heading “tail first”.

### 2- Pinned +1/2 defects

Since there was no creation of new defects after confluence, annihilation events involving pairs of defects of opposite charges resulted in a decrease of the surface density of +1/2 defects with time as previously reported (26) (Figure 1d). However, this density did not completely vanish but plateaued at 1.8 ± 0.3 defects/mm^2^, ^~^ 5 h before t_2L_. Strikingly, these remaining defects remained fully immobile (“pinned”) within the monolayer (Supplementary Figure 3). Tracking them backwards in time showed that they actually kept their positions up to 15 h before the transition to bilayering. This non-motile behavior was not caused by a global jamming of the monolayer (41) as individual cells still moved along the lines of the orientation field (Supplementary Movie 1). Cells located in the defect’s head converged toward its core. In contrast, no significant flow was detected in the tail region. As a result, the average velocity profile along the axis of an average +1/2 immobile defect was very asymmetric about the core with non-zero values only at the head (Figure 1e). This observation is qualitatively reminiscent of the asymmetric flow field around defects in monolayers of Neural Progenitor Cells (8) although the NPC system is extensile and therefore, in that system, the flow is directed from the tail toward the defect’s core and decreases sharply at its head. In the present contractile case, the flow distribution is opposite with measurable flows only at the head of the defect. In addition, we observed an accumulation of C2C12 cells at the head of the pinned +1/2 defects (Figure 1f) which preceded the formation of 3D structures. Analogy with ref (8) then strongly suggested that this 2D cell accumulation participated to the formation of 3D structures.

To access the force field at these motionless +1/2 defects, we took advantage of the high sensitivity of Focal Adhesions (FAs) to an external force (42, 43). FAs were imaged via the labelling of paxillin. Close to the core of the defects, they were classically positioned at the end of the stress fibers (Figure 2a) but their area and shape were not uniformly distributed: The cells of the tail close to the core had larger FAs elongated in the tail direction (Figure 2b-d). This observation is consistent with a force acting on the tail cells, preventing the motion of the +1/2 defects.

**Figure 2:**
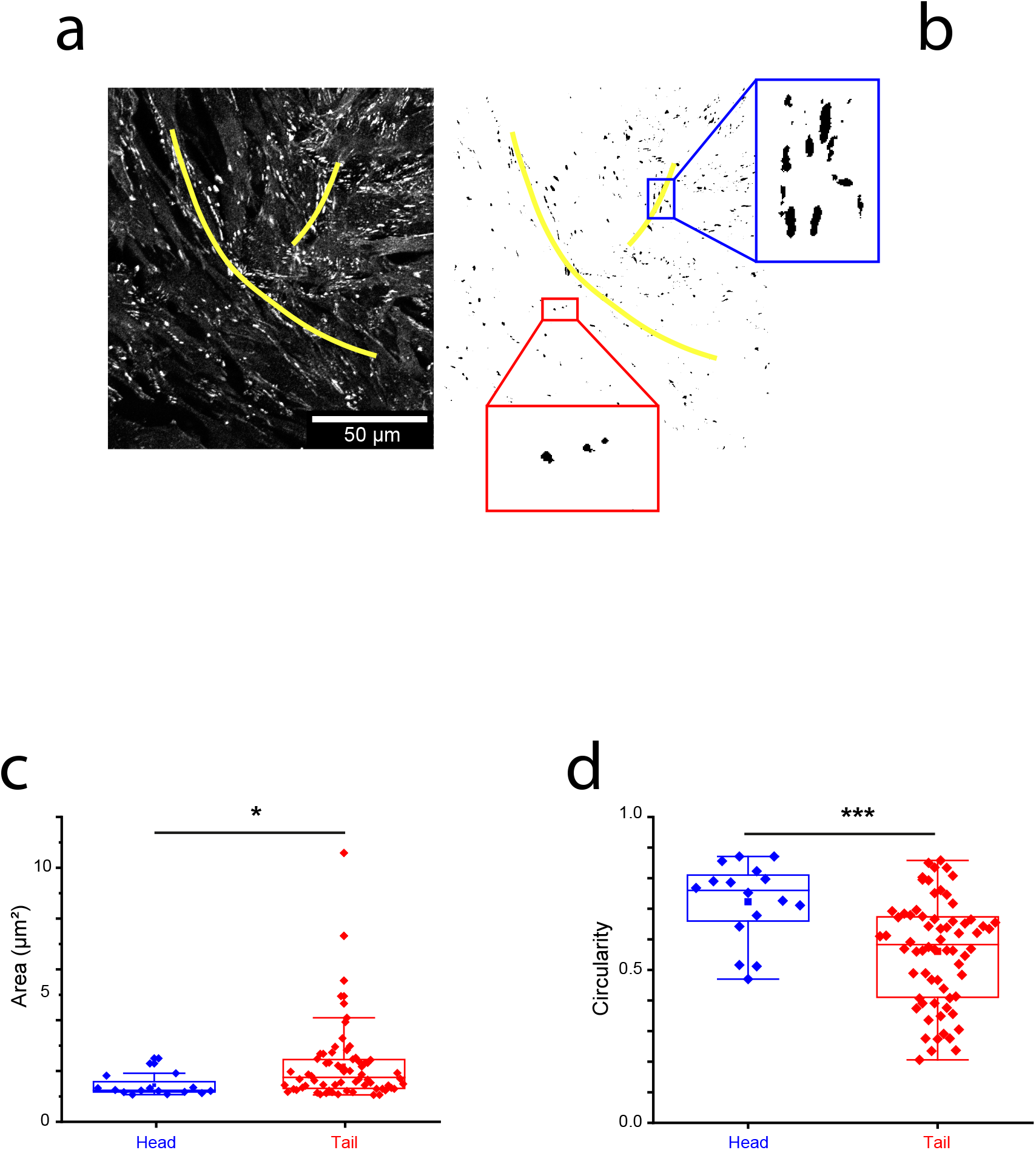
a/ Focal adhesions (FAs) (fixed cells, paxillin labeling) at an immobile defect just before bilayering. The defect has been outlined in yellow. b/ Thresholded image. Note the elongated large FAs in the tail while they are smaller and more circular in the head. c,d/ Quantification of the area and circularity of the FAs in the tail and head regions of the immobile defects (5 independent defects). FAs in the tail are elongated in the direction of the defect.

Since +1/2 topological defects have been previously shown to play an important role in apical cell extrusion processes (8, 17), it was interesting to question how the transition of C2C12 cells from a monolayer to a bilayer could be related to this particular cell organization and to the associated velocity and force fields.

### 3- Onset of bilayering at pinned +1/2 defects

Previous reports have emphasized the importance of +1/2 defects in the formation of 3D disorganized clusters from a monolayer (8). However, in contrast with these previous studies, the transition evidenced here is characterized by a coordinated bilayering process and not by the extrusion of a disorganized cell aggregate.

In the present situation, the 2D-3D transition occurred exclusively at the motionless +1/2 defects via the splitting of the monolayer in two stacked layers. Further evolution of the system was driven by the migration of these two layers. Indeed, we observed that cells initially at the head of the defect but close to its core, collectively migrated *on top* of the cells of the first layer pertaining to the tail of the initial defect. In parallel, the cells initially in the core vicinity but pertaining to the tail of the defect, moved collectively *under* the cells originally at the head of the defect (Figure 3, Supplementary Movie 2).

**Figure 3:**
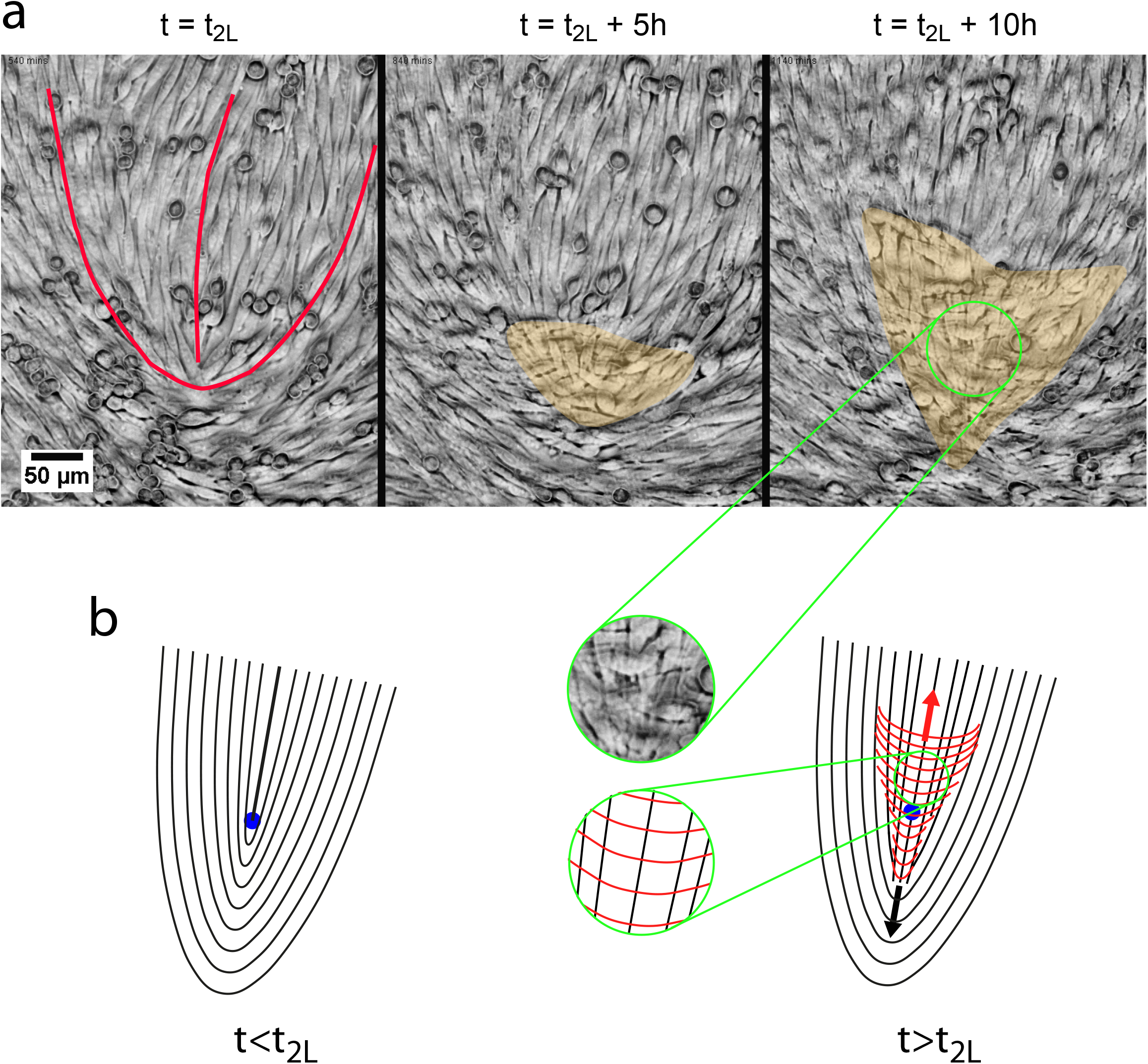
a/ Sequence of snapshots (phase contrast) illustrating the progression of the bilayering. The orange area highlights the crisscross bilayer. The outline of the defect at t=t_2L_ is shown as a red line. See full movie in Supplementary Movie 2. b/ Schematic of the bilayering process. The red lines correspond to the orientation field lines of layer 2 while the black lines refer to layer 1. The blue dot shows the initial position of the core. Note the crisscross organization of the two layers when t>t_2L_ (insets).

It is worth noticing here that, more than 50 years ago, by carefully observing sheets of embryonic lung fibroblasts, T. Elsdale and co-workers concluded that crisscross bilayering originated at immobile “ridges” between well-oriented domains (33). Given the technology of the time, this was a truly remarkable conclusion. Here, we largely confirm this part of Elsdale’s analysis, and we now identify these ridges with pinned +1/2 nematic singularities.

### 4- Organization and dynamics of the bilayer

At the onset of bilayering at a pinned +1/2 defect, the cells of layer 2 originated from the head of the initial defect and migrated on top of layer 1. Similarly, the cells of layer 1 that crawled under layer 2 originated from the tail of the initial defect. Since cells of both layers kept their initial orientations during this process, the cells of layer 1 and layer 2 were oriented along perpendicular directions with respect to each otherin the region of overlap. These orthogonal orientations therefore correspond to the previous observations of fibroblast-like cells from lung (44) or heart (18). In the following, we call this process “crisscross” or “perpendicular” bilayering.

Perpendicular bilayering was not the only way by which the thickness of the cell sheet increased: In defect-free regions, cells dividing in the confluent monolayer could not intercalate in the monolayer and adhere to the substrate due to lack of space. They partially incorporated in layer 1 while keeping their initial orientation, contributing therefore to the progressive development of layer 2. Therefore, this configuration yielded a gradual increase of thickness in which cells pertaining to both layer were all oriented along the same direction. We call this process “parallel bilayering”. We emphasize that, whereas perpendicular bilayering is clearly a collective event implying the concerted action of the cells in the above-mentioned antiparallel displacements, parallel bilayering is an individual insertion event originating from cell division. Although both mechanisms contribute to the multilayering of this cell system, they are clearly independent.

We now turn to the dynamics of the perpendicular bilayering. The migration modes of layers 1 and 2 were primarily collective with only rare events of cells detaching from their neighbors at the respective front edges. On the axis of the initial defect, the front edge velocities of layers 1 and 2 with respect to the glass substrate measured by tracking were comparable (respectively 12±4 μm/h (n=7) and 15±7 μm/h (n=5)). Recirculating flows were observed on the sides (Supplementary Movie 1). Addition of EDTA that disrupts the cadherin-cadherin interactions did not abrogate perpendicular bilayering (3 independent experiments) (Supplementary Figure 4).

Importantly, the direction of collective migration was parallel to the cells’ main axis for layer 1 but orthogonal to it for the cells of layer 2 (Supplementary Movies 1,2). Note that isolated C2C12 cells migrate in the direction of their long axis.

We conclude that perpendicular bilayering initiates exclusively at +1/2 immobile defects as a result of the accumulation of cells at the defect head. The monolayer then splits into two stacked layers that migrate actively in antiparallel directions on top of each other. Because they keep their initial respective orientations in this process, the cells of the two layers end up being perpendicularly oriented.

### 5- Extra Cellular Matrix secretion at apical side mediates multilayering

The migration of a second layer on top of the first one questions the biochemical nature of the surface on which cells of layer 2 collectively migrate (i.e. the apical side of layer 1). In particular, extrusion of MDCK or NPC cells did not lead to a well-developed second layer but to aggregates or apoptotic cells.

Since cadherins appear not to be involved in this process, we hypothesized that interactions between successive layers were mediated by cell-secreted ECM proteins and we therefore proceeded to precisely localize some of its prototypic components in the stacked layers.

We focused on 3 major proteins of the extracellular matrix (45): i/ collagen IV, which is the structural element of ECM, ii/ fibronectin, and iii/ laminin, that both directly interact with integrins (35, 46). Information on cell anchoring in these multilayered systems was obtained in parallel by imaging paxillin that is part of the Focal Adhesion complex (47, 48).

Cells were fixed at different stages of multilayering and the proteins of interest were imaged by confocal microscopy. On well-formed multilayers, we systematically observed the presence of the three ECM proteins sandwiched between successive stacked layers (Figure 4a-c). Paxillin imaging showed Focal Adhesions for cells of all layers (Figure 4a-e), confirming that cells of superimposed layers interacted with ECM via integrin-mediated adhesions. Therefore, it appears that cells secrete the ECM molecules that are necessary to ensure good adhesion and friction between the successive stacked cell layers.

**Figure 4:**
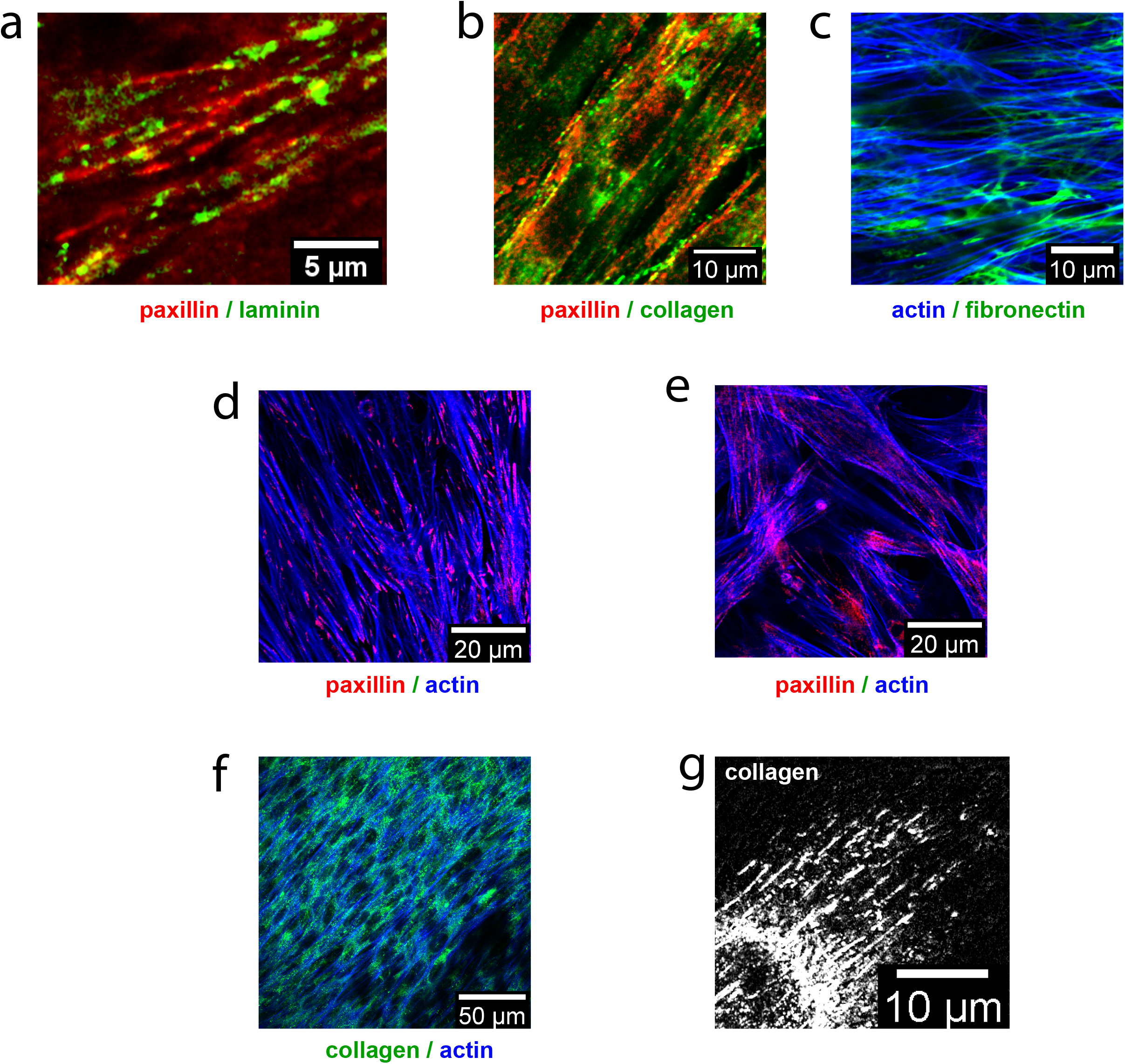
a-c/ Laminin, collagen and fibronectin can be found between the cells of layers 1 and 2 in a multilayered sample (these images are confocal slices acquired 10 μm above the glass surface). Note the colocalization of ECM proteins with the Focal Adhesions, characteristic of the 3D organization of the cells. d,e/ The FAs structure is very different for layer 1 interacting with glass (d) compared with the subsequent layers that interact with the ECM that is produced by the underlying cells (e). The fibrillar structure of the FAs that colocalises with actin within the multilayered assembly is typical of a tridimensional organization of the cells. (confocal slice on glass (d) and 10 μm above the glass surface (e). Fixed cells). f/ ECM (here collagen) is detected at the apical surface of the topmost layer and follows the direction of the cells. g/ At the top most apical surface, collagen organizes itself in the form of parallel fibers in the direction of elongation of the cells forming submicron guiding “rails” (super-resolution image).

Focal adhesions looked similar for all layers except for layer 1 that was in direct contact with the Fibronectin-coated glass while the other layers interacted with the apical side of the underlying cell layer (Figure 4d,e). Except very close to the defect core at the mono- to bilayer transition, the adhesion sites of layer 1 on glass were well-defined Focal Adhesions at the extremities of stress fibers as it is usually observed for cells spreading on ECM coated glass (Figure 4d). In the subsequent layers, the adhesions that the cells developed with their underlying cell layer were more fibrillar (Figure 4e) and colocalized with the ECM with a pattern typical of 3D architectures (Figure 4a,b) (47, 49). The adhesions of the cells of layer 1 with its glass substrate were therefore different from the ones developed between the stacked layers.

Furthermore, whatever the number of layers, we systematically detected all 3 ECM proteins at the apical side of the *topmost* layer (for instance on top of layer 1, before layer 2 had started developing) (Figure 4f). Adding low-concentration collagenase (that destroys collagen) just before t_2L_ totally prevented the development of layer 2. The cells remained in the form of a monolayer for at least 10 h after the addition of the drug. The enzyme also destroyed the limited patches of layer 2 that had already begun to form (Supplementary Movie 3). The relative low impact of collagenase on the adhesion of layer 1 to glass confirms our previous observations that Focal Adhesions for layer 1 are different from layer 2 and the others.

From these observations, we conclude that the cells of the topmost layer secrete a collagen IV-rich ECM layer at their apical side on which the next layer migrates in the perpendicular multilayering process. Consistently with this observation, we couldn’t detect paxillin at the topmost apical surface.

Interestingly, the collagen secreted at the topmost apical layer was not uniformly distributed but took the form of well-oriented parallel bundles of width ^~^ 500 nm, separated by ^~^ 1000 nm (Figure 4g). These bundles were oriented in the direction of the cells that secreted them. Since cells of the head of the comet migrate on top of layer 1 in the direction given by the cell bodies of this first layer (i.e. the direction of the tail), this collective motion may have been mediated by the structured ECM secreted by these cells at their apical side. During this process, the layer 2 migrating cells didn’t reorient and therefore migrated perpendicularly to their long axis. Of note, layer 1 migrated actively under layer 2 along the direction of its constituent cells (therefore perpendicular to the cell orientation of layer 2).

## Discussion

Cultures of C2C12 cells shortly after confluence show the characteristic flow field of an active nematic contractile system around +1/2 defects. However, a fraction of these defects remains immobile in the 15 h preceding the bilayering. These stationary defects eventually give rise to perpendicular bilayering events and, vice-versa, all perpendicular bilayering events originate at these defects.

In the case of extensile systems, a skewed velocity distribution has previously been attributed to an “anisotropic friction” that takes larger values when the velocity of the cells is perpendicular to their orientation (8, 50). For extensile systems, this velocity profile results in an accumulation of cells that eventually extrude from the monolayer.

In the present case, at high cell density, some of the defects get pinned down on the surface. Recent work has estimated the force necessary to stop a defect (51). Here, the arrest of these defects can be mediated by heterogeneities in the basal deposition of ECM by the cells on the glass surface (52, 53) (Supplementary Figure 5). Once the core is pinned down, the flows at the defect are considerably perturbed. Along the axis of the defect, the only subsiding displacements are at the head, directed toward the core. Interestingly, these flows are perpendicular to the cells’ long axes. The tail cells then resist statically to the flow-induced force as can be evidenced directly by the shape and orientation of the FAs (42).

When the cell density becomes too large, the force exerted by the head cells becomes too high and the system doesn’t remain bidimensional. An out-of-plane force develops and the head cells cross over to form a layer 2 migrating on the apical side of layer 1, while layer 1 migrates under the head cells in the opposite direction.

Migration of layer 2 on top of layer 1 is critically dependent on the secretion of ECM proteins on the apical side of layer 1 as demonstrated by the inhibition of formation of a bilayer in presence of collagenase. Indeed, MDCK epithelial cells that don’t secrete these proteins on their apical side (54) don’t provide the adequate template for another layer to spread on them and therefore don’t form bilayers. Rather, MDCK cells at the defect sites individually extrude but can’t adhere on the apical side of the monolayer and die (17, 55, 56). Since disrupting N-cadherins-mediated C2C12 cell-cell contacts with EDTA did not abolish the bilayering process, the role of the cadherins in these migrations appears to be secondary compared to the cell-ECM interactions.

Not only are ECM proteins abundant at the apical surface of the topmost layer but they are also structured in the direction of the cell bodies as well-defined sub-micron bundles separated by typically 1 μm in the direction of the cells that have produced them. Our observations suggest that these “tracks” may guide the layer migrating on them. Indeed, microgrooved substrates of subcellular periodicity have been shown to guide the migration of cells or cell assemblies by Contact Guidance (57–60).

Interestingly, cells migrating on top of layer 1 to form layer 2 keep their initial orientation, meaning that their velocity is perpendicular to their long axis. This mode of “sideways” migration in collective migration, although uncommon, has occasionally been observed for instance at the front edge of a migrating monolayer of Zebra Fish epicardial cells (61) or at the edge of closing wounds in fibroblasts’ monolayers (62). Since some features of the cells such as the morphology of their FAs bears some characteristics of a 3D environment, they may also have switched to a mode of migration intermediate between 2D and 3D. Of notice, C2C12 cells have been reported to orient perpendicularly to castellated sub-micron ridges whose dimensions are very close to the collagen bundles that develop at the apical side (63).

The speed of the collective migration after the initiation of the bilayering was measured to be ^~^15 μm/h for both cell layers (in both cases relative to the glass substrate). The velocity of layer 1 is within the range of values classically measured in wound-healing experiments. However, cells forming layer 2 collectively migrate on a counter-migrating substrate (the layer 1). They run the wrong way on a treadmill at a relative large speed of ca. 30 μm/h. At these large speeds, the friction between these two layers can also contribute to maintain the cells of layer 2 perpendicular to their velocities by preventing cell reorientation.

At the end of this process, the structure adopted by the cells is a perpendicular crisscross orientation between cells of layers 1 and 2.

Altogether, this study highlights the complex interplay between physical and biochemical cues resulting in the crisscross organization of stacked layers of muscle cells. We propose that such mechanisms may be at play in vivo at the onset of the development of crisscross muscle layers in hydrostats-like structures such as the small intestine (22) or the developing hydra where the endoderm and the ectoderm have perpendicular orientations (20, 21).

## Methods

### Cell Culture and Plating

The experiments were conducted with (i) wild type C2C12 cells (gift from Clotilde Thery (Exosomes and Tumor Growth, INSERMU932, Institut Curie)), (ii) CRISPR labelled Actin M-cherry cells and (iii) LifeactGFP cells that were generated and characterized in-house in the BMBC facility.

Cells were cultured in standard DMEM medium (Thermofisher scientific) supplemented with 10% FBS (Gibco) and 1% Pen-Strep Penicillin-Streptomycin mixture (Gibco). Experiments involving fluorescence microscopy of any kind were performed with phenol red free Gibco™ DMEM, high glucose, HEPES, supplemented with 10% FBS (Gibco) and 1% Penicillin-Streptomycin(Pen-Strep) mixture (Gibco) and Glutamax supplement.

For all experiments, cells were plated on Fibronectin-coated (10 μg / mL) glass coverslips. Note that, due to their large passage number after reaching confluence (more than 30), the cells have lost their capability to differentiate.

The concentration of EDTA used for calcium chelation to disrupt N-cadherins was 2.5 mM. The concentration of collagenase used was 2.5 microunits/mL.

### Microscopy

The phase-contrast and epifluorescence time lapse movies on live cells were acquired with an automated Olympus X71 microscope, equipped with 4x, 10x and 20x objectives while maintaining 37°C, 5% CO2 partial pressure and 95 % relative humidity (Life Science incubator).

The acquisitions were performed with a CCD camera (Retiga 4000R, QImaging) controlled by Metamorph (Universal Imaging). The typical frequency of image acquisitions was 15 minutes.

For confocal time-lapse imaging, we used an Inverted laser scanning confocal microscope with spectral detection (LSM700 - Zeiss) equipped with a CO2 incubator to observe fluorescent live cells with a 25x oil objective. The typical duration between acquisitions was 30 minutes.

For super-resolution confocal imaging on fixed cells, we used an inverted Laser Scanning Confocal Microscope with Spectral Detection and Multi-photon Laser (LSM880NLO/MaiTai Laser - Zeiss/Spectra Physics), 63x oil immersion objective combined with super-resolution technique Airyscan.

Tracking experiments were performed by using mixtures with 10% Crispr labelled Actin M-cherry cells. Observation were performed in epifluorescence. Thanks to the relatively large depth of field, cells of both layers could be imaged simultaneously and then tracked manually.

### Cell Fixation and Fluorescent Staining

To fix the cells, a 16 minutes of 4% (wt/vol) PFA treatment was followed by permeabilization by 0.1% Tween-20 (10 mins). Fibronectin staining was an exception, for which ice-cold 100% methanol for fixation (no permeabilization step) was used. DAPI was used for the nuclei, phalloidin-Tritc and SIR-Actin dye were used for actin and respective primary and secondary antibodies were used for respective proteins (see Supplementary Table 1 for details). For live cell imaging, SIR actin and SIR DNA were used.

In Figure 1a, the cells were fixed in ice cold ethanol and acidic Harris Hematoxilin solution (HHS16, Sigma) (33).

### Analysis of the images

Analysis of the images was performed using FIJI (64) or Matlab (MatWorks).

For the analysis of the Focal Adhesions (Figure 2), paxillin was imaged and the images were thresholded. FAs smaller than 1μm^2^ were discarded. The remaining FAs were analyzed for their area and circularity defined as 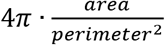.

The density of cells was mapped by manually counting the nuclei (SIR Nucleus labelled cells).

### Orientation and Velocity Field Extraction

The orientation field was acquired by computing the local gradient structure tensor with the FIJI ‘OrientationJ’ plugin, as described in (26, 65). To map the velocity fields, Particle Image Velocimetry (PIV) (39) was performed with the Matlab toolbox PIVlab 1.41 (66) with 16 pixels-windows, which corresponds to 29.7 μm, and 50% overlap. The time intervals between successive frames were 15 minutes or 30 minutes depending on the experiment.

### Defect detection and Analysis

Defects were detected by mapping a local order parameter in windows of size *Ω* = 27.8 μm * 27.8 μm:

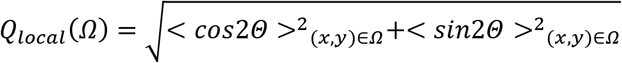

whose minima correspond to the position of the defects. Their charge and orientation was measured by calculating the variations of angle along a virtual loop around the core (28).

The extracted orientation and velocity fields at +1/2 defects were then processed to massively average these two fields (typically, over 5400 acquisitions from 350 independent defects that have been aligned together).

## Acknowledgements

It’s a pleasure to thank F. Brochard, JL Maitre, O Ozguc, J. Prost and the members of the “Biology-inspired Physics at MesoScales” (BiPMS) group for discussion and advice.

Trinish Sarkar PhD was funded by the IC3iPHD program, which has received funding from the European Union’s Horizon 2020 research and innovation program under the Marie Skłodowska-Curie grant agreement No 666003.

Victor Yashunsky gratefully acknowledges the Cell(n)Scales Labex and the European PRESTIGE program (PCOFUND-GA-2013-609102 grant).

We thank the Cell and Tissue Imaging core facility (PICT IBiSA), Institut Curie, member of the French National Research Infrastructure France-BioImaging (ANR10-INBS-04) and, in particular, Olivier Renaud and Olivier Leroy.

We thank the BMBC facility of the PCC laboratory for guidance and help and, in particular, Aude Battistella for the CRISPR insertion, Fanny Cayrac for the Lifeact-GFP plasmid insertion and John Manzi for biochemical analyses.

The BiPMS group is a member of the Cell(n)Scales Labex (grants ANR-11-LABX-0038, ANR-10-IDEX-0001-02) and an associate member of the IPGG.

## Figure captions

**<u>Supplementary Figure 1:</u>** Confocal xz section of a fixed mature multilayered cell sheet, t=3 days. The actin and the nuclei are labelled (actin blue, nuclei red). The stack consists of four to five cell sheets on top of each other. Its thickness goes up to 30 microns.

**<u>Supplementary Figure 2:</u>** a/ C2C12 cells at confluence organize themselves in well-aligned domains between which topological defects position themselves. The color codes for the orientation. b/ −1/2 defect (actin labeling), c/ +1/2 defect (actin labeling). Red lines outline the defects. d/ Schematic of a +1/2 comet-like defect and notations used in the article.

**<u>Supplementary Figure 3:</u>** Representative trajectories of an immobile and a mobile +1/2 defect over a time course of 5 h. Mobile defects behave as self-propelled particles while immobile defects remain in place within our experimental accuracy.

**<u>Supplementary Figure 4:</u>** Crisscross bilayer in presence of EDTA (live cells, phase contrast, 20 hours post confluency). EDTA (10mM) was introduced in the medium after cells have reached confluency. Although it impairs cadherin-mediated cell-cell adhesions, EDTA does not abrogate the crisscross structure (see inset where the orientation is color-coded and the typical orthogonal structure is visible)

**<u>Supplementary Figure 5</u>** Secreted ECM at the glass surface, along with the basal side of layer 1 (green: collagen, red: actin). Mature stack (day 3). Note the inhomogeneous distribution of the collagen. Fixed cells.

**<u>Supplementary Movie 1:</u>**

http://xfer.curie.fr/get/0ceKBvP8C2a/supp%20movie%202%20tracking%20of%20cells.avi

Before the bilayering and during this process, complex flows can be visualized on the sides of the comet on top of the global progression of the cells forming layer 2. For this movie, 10% of the cells were MCherry Crispr-labeled so they can be tracked independently.

**<u>Supplementary Movie 2 :</u>**

http://xfer.curie.fr/get/ChGUA58A8sI/supp%20movie%201_phase.avi

Progression of the bilayering process with time. Phase contrast movie. Note the crisscross orientation of the bilayer. See corresponding sequence in Figure 3.

**<u>Supplementary Movie 3</u>**

http://xfer.curie.fr/get/mRK6kBVTtFp/supp%20movie%203%2019_12_18_collagenase_C2C12_mixed-5.avi

Addition of collagenase impairs bilayering and destroys the partial bilayers that have started to form. Collagenase was added at the onset of bilayering (immediately before the start of this video).

**<u>Supplementary Table 1</u>**

Reactants and concentrations used in the present work.

