## Supplementary material for "Crisscross multilayering of cell sheets": supp fig 1-5

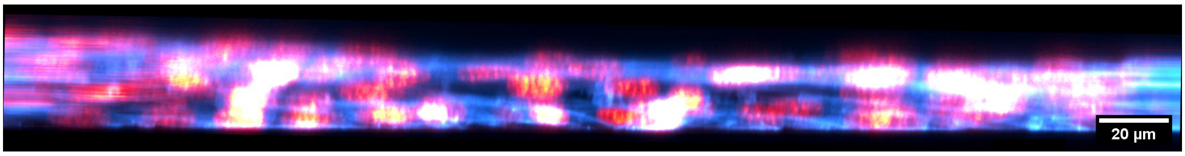


### Supplementary Figure 1

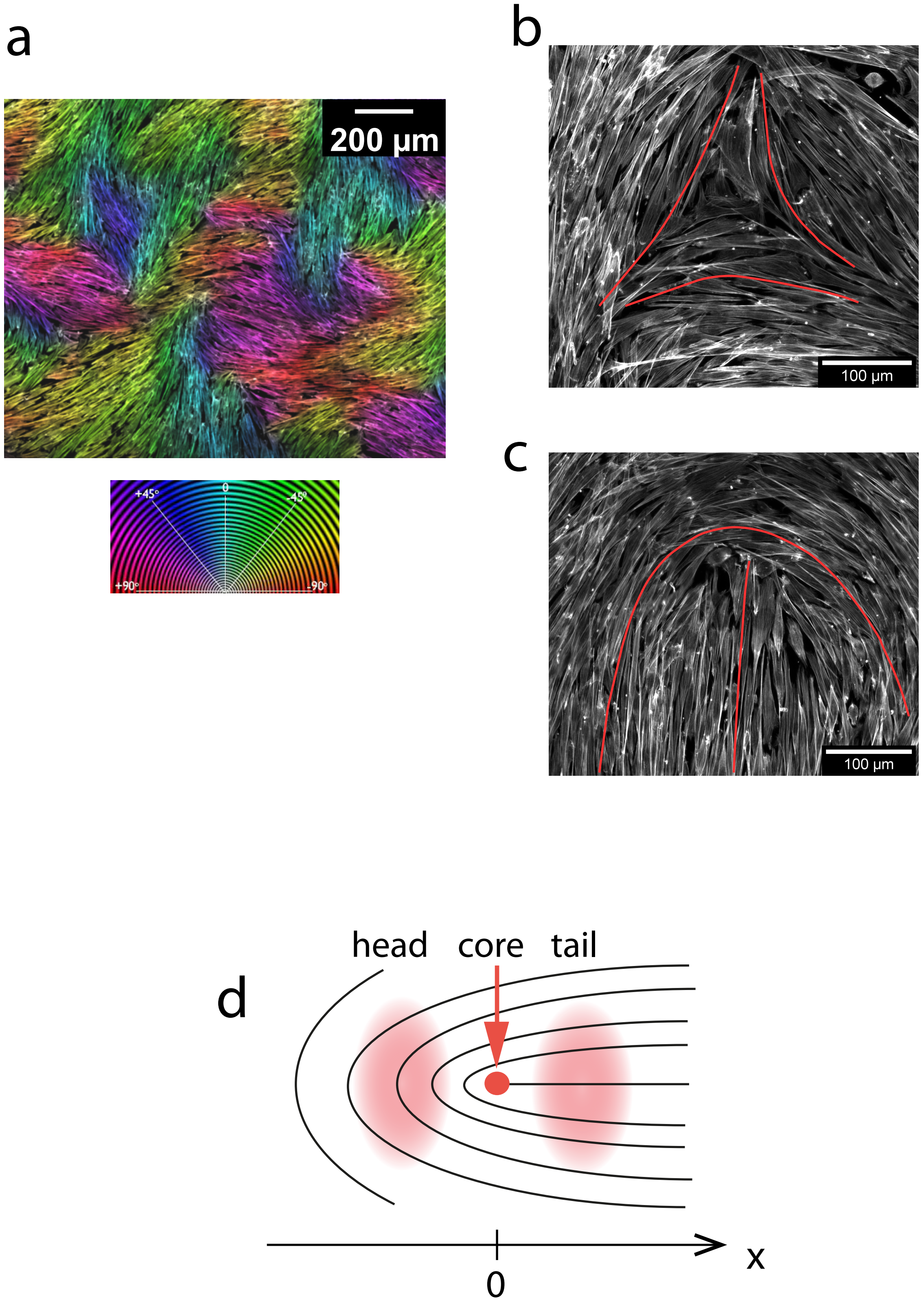


Supplementary Figure 2

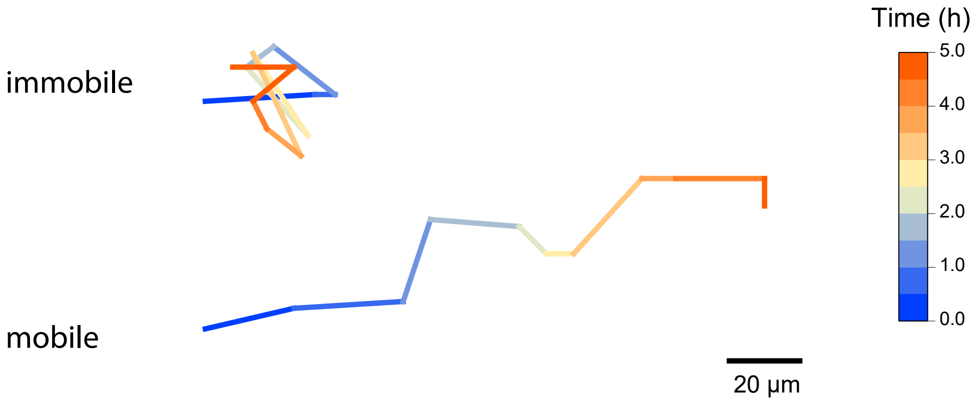


### Supplementary Figure 3

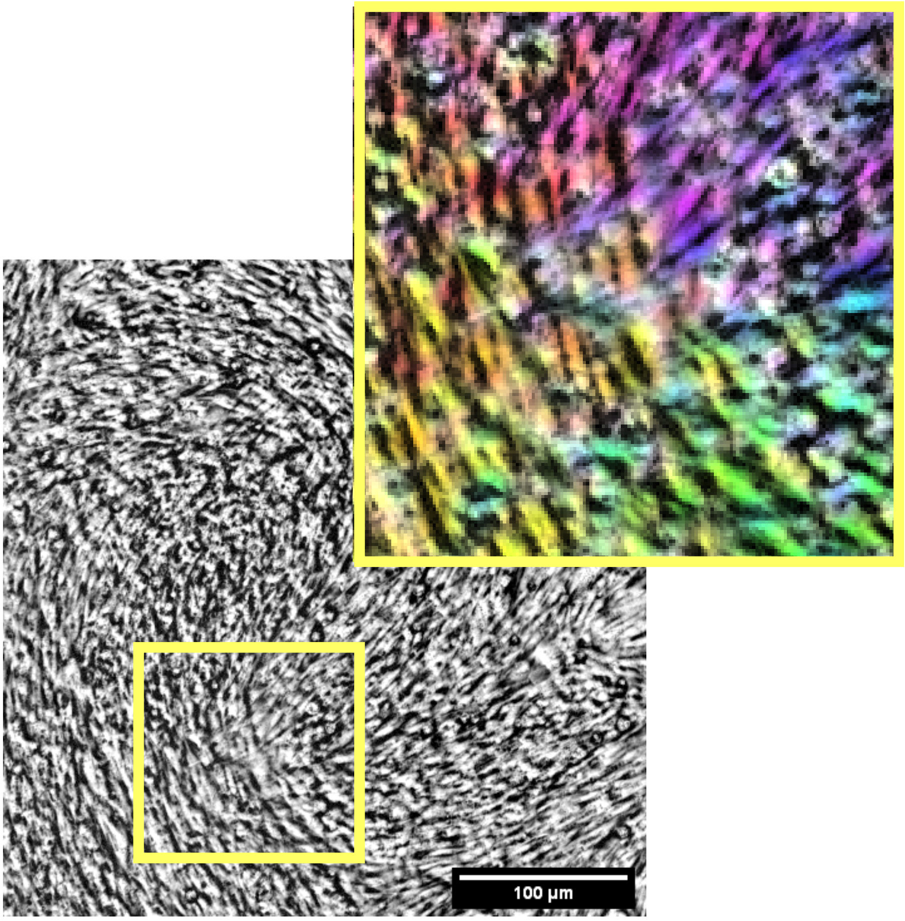


### Supplementary Figure 4

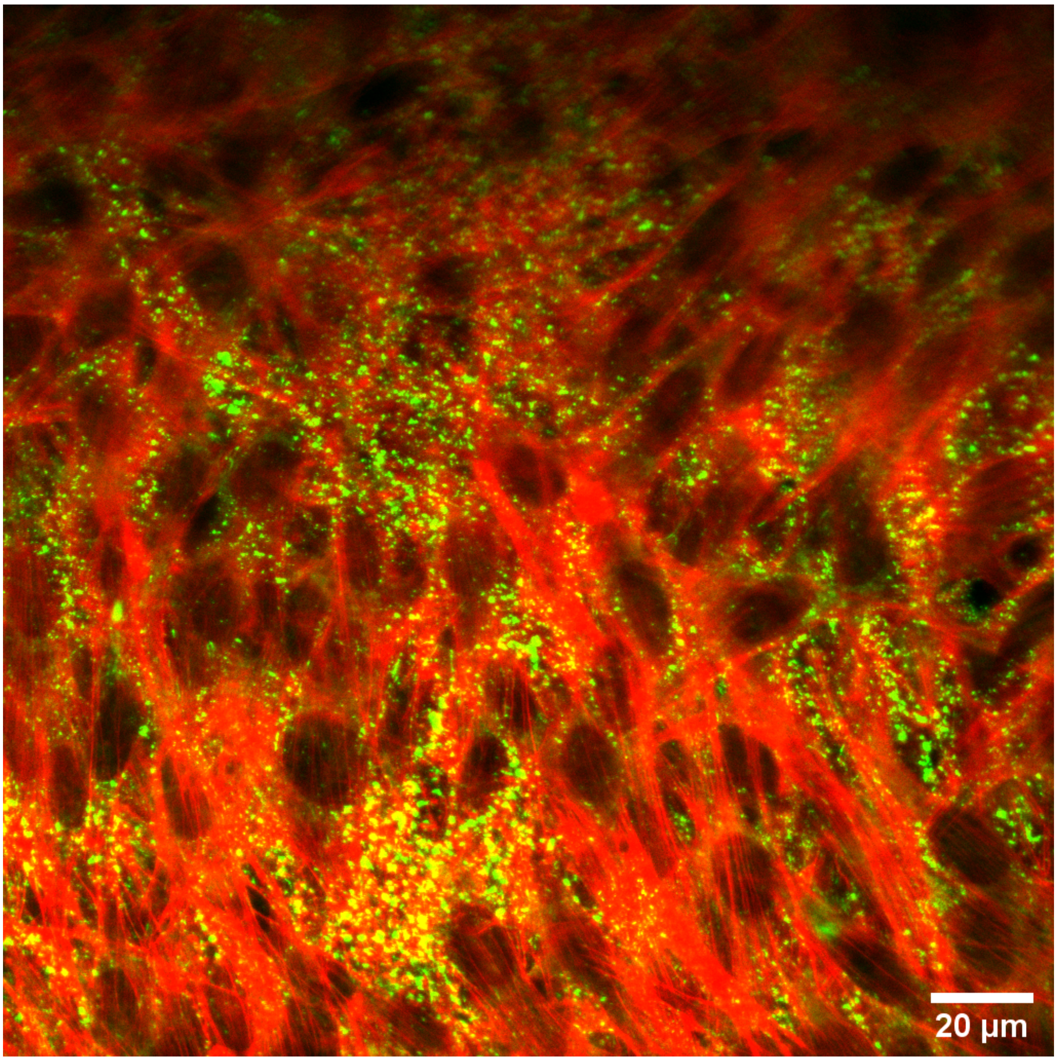


Supplementary Figure 5

| Target | Type | Reference | Dilution |
|---|---|---|---|
| Laminin α1 and α2 chains | Rabbit Laminin poly 1+2 | ab7463(abcam) | 1:100 |
| Paxillin | Rabbit mono(with Alexa Fluor®647) | ab246719(abcam) | 1:50 |
| Collagen IV | Rabbit polyclonal to Collagen IV | ab6586(abcam) | 1:100 |
| N-cadherin | Mouse Monoclonal | 33-3900(Thermofisher) | 1:50 |
| Actin | Phalloidin-TRITC | P1951 (Sigma-Aldrich) | 10 μM |
| Actin | Sir-actin | Tebubio(SC001) | 0.1 μM |
| Fibronectin | Rabbit polyclonal to fibronectin | F3648 (Sigma) | 1:100 |

| Name | Type | Dilution |
|---|---|---|
| Mouse Secondary Antibody | GoatXMouse Alexa Fluor PLUS 647(A32728) ( Thermofisher) | 1:200-1:300 |
| Rabbit Secondary Antibody | ChickenXRabbit Alexa Fluor 488(A-21441) (Thermofisher) | 1:100 |

### Supplementary table 1
